# Brain Signal Variability During Rest as a Neural Mechanism Underlying Cognitive Reserve

**DOI:** 10.1101/2025.04.09.648011

**Authors:** Annabell Coors, Yaakov Stern, Christian Habeck

**Affiliations:** Cognitive Neuroscience Division, Department of Neurology, Columbia University Vagelos College of Physicians and Surgeons, New York, NY, USA; Department of Psychiatry, Psychotherapy and Psychosomatic Medicine, University Medical Center Halle, Halle (Saale), Germany; The Taub Institute for Research in Alzheimer’s Disease and the Aging Brain, Columbia University, New York, NY, USA; The Gertrude H. Sergievsky Center, Columbia University, New York, NY, USA

**Keywords:** resting-state fMRI, cognition, resilience, aging, brain reserve

## Abstract

**Background:** Resting-state brain signal variability has been found to vary with age and cognitive function. Neural flexibility has been suggested as a neural mechanism underlying cognitive reserve (CR), a construct that describes better than expected cognition given brain status. Thus, we examined the associations between age, resting-state brain signal variability, cognition, and CR.

**Method:** Analysis was based on resting-state functional neuroimaging data from 470 participants (aged 20-80 years) from the Reference Ability Neural Networks and the CR studies. Brain signal variability was quantified for each brain region as the log-transformed standard deviation of the time-varying blood-oxygen-dependent (BOLD) signal. We then derived variability patterns related to age, perceptual speed, fluid reasoning, episodic memory, and vocabulary using Scaled Subprofile Modelling principal component analysis. To perform the formal test whether these patterns fulfill the requirements for CR, we examined whether they explained additional variance in cognition beyond brain status, age, sex, and education, or moderated the brain status-cognition relationship. We additionally stratified all regression models by age (cutoff: 60 years) and sex.

**Results:** BOLD signal variability showed an age-related increase in subcortical/medial brain regions, and an age-related decrease in cortical regions. It also met the CR test for speed (standardized regression coefficient (ß)=0.251, 95% confidence interval (CI): 0.118-0.384, *p*_FDR_<0.001), episodic memory (ß=0.344, CI: 0.200-0.489, *p*_FDR_<0.001), reasoning (ß=0.316, CI: 0.197-0.436, *p*_FDR_<0.001), and vocabulary (ß=0.270, CI: 0.167-0.373, *p*_FDR_<0.001). Associations were stronger in women for vocabulary and in young individuals for reasoning.

**Conclusions:** BOLD signal variability plays a role in aging and cognition and underlies CR.

## 1. Background

Intra-individual variability in brain activity, as measured by the deviation of a brain signal over time, has often been overlooked or considered “noise”. However, previous studies suggest that it represents an informative dimension about the brain’s functional integrity and flexibility that is not captured by the mean-based signals.^1–4^ While greater intra-individual variability in brain activity may imply greater flexibility due to more exploration of available functional network configurations,^2,5–7^ it has also been found that temporal variability is lower during task activation than during rest, which may allow for better signal transmission.^8,9^ Further, intra-individual variability in brain activation of selected regions during rest has been suggested as a marker of various diseases, including functional neurological disorders^10^ and Alzheimer’s disease.^11^ However, before evaluating its potential as a disease-related biomarker, it is crucial to first understand the associations between age and intra-individual brain signal variability, as well as its relevance for cognitive performance.

Regarding the association between age and brain signal variability, a resting-state functional magnetic resonance imaging (rs-fMRI) study of 83 children aged 2 to 8 years found that older age was associated with higher blood-oxygen-level-dependent (BOLD) signal variability in the frontal, temporal, and parietal regions and lower BOLD signal variability in more restricted regions and medial structures, such as the hippocampus and the parahippocampal gyrus.^12^ An electroencephalography (EEG) study on differences in brain signal variability during a face-recognition task between children aged 8 to 15 years and young adults aged 20 to 33 years, found that brain signal variability was higher in the group of young adults compared to the children.^13^

In terms of the development of brain signal variability from young adulthood to older age, an rs-fMRI study of 19 adults aged 20 to 30 years and 28 adults aged 56 to 85 years reported lower overall brain signal variability in older adults.^4^ However, this study also found strong regional differences in brain variability. While some regions belonging to the default mode network (e.g., angular gyrus, middle temporal gyrus, and superior medial gyrus) and other areas relevant to motor control and sensorimotor processing (e.g., postcentral gyrus, Rolandic operculum, and supplementary motor area) showed lower BOLD signal variability in the older adult group, variability was higher in the older adult group in the superior frontal gyrus, the inferior temporal gyrus, and the cerebellum.^4^ Age-related increases in BOLD signal variability in the superior frontal gyrus, the inferior temporal gyrus, and the cerebellum have also been reported in another rs-fMRI study^14^ and one task-related fMRI study.^15^ Further, the task-related fMRI study also found age-related decreases in brain signal variability in some regions of the default mode network (e.g. angular gyrus, middle temporal gyrus, superior frontal gyrus) and other areas relevant to motor control, sensorimotor processing, and visual processing (precentral gyrus, postcentral gyrus, calcarine gyrus).^15^ Region-specific age-related differences in brain variability have also been reported in a rs-fMRI study from the Nathan Kline Institute (NKI-Study) of 378 participants that spanned the wide age range of 6 to 85 years. However, in this study, those brain regions that included a key node of the salience network tended to have larger variability in brain activation in older individuals, while variability in most other regions tended to be lower in older individuals.^16^ Another rs-fMRI study, which included a group of 20 cognitively normal adults and a group of 20 cognitively impaired individuals, found that older age was associated with higher voxel-wise calculated brain signal variability only in the cognitively impaired group.^17^

Studies on the role of brain signal variability in cognitive performance have shown that brain signal variability appears to be associated with cognitive function, but the direction of this association across cognitive domains would benefit from further investigation. For example, the developmental trend toward greater EEG signal variability from childhood to young adulthood was found to be accompanied by an increase in face-recognition accuracy and a decrease in reaction time variability, but it was unrelated to mean reaction time.^13^ This was interpreted as brain signal variability being mainly relevant for higher behavioral consistency.^13^ In rs-fMRI studies that included adults in the age range of 19 to 87 years, higher BOLD signal variability has been found to be associated with better fluid intelligence in some,^18,19^ but not all studies.^20,21^ Findings on the association between rs-fMRI BOLD signal variability and perceptual speed are also mixed, with one study reporting that higher BOLD signal variability is associated with speed for only one of two time points,^19^ and another study reporting no association.^18^ There may also be regional differences, as one study showed that higher variability in most, but not all brain regions, was associated with faster speed.^15^ Regions where higher variability was associated with slower speed included the cerebellum and subcortical structures (e.g., the right caudate nucleus and hippocampus).^15^ Results are also mixed for various aspects of memory that have been studied.^17–19^ A rs-fMRI study of 91 healthy adults aged 60 to 80 years found that greater voxel-wise brain signal variability in motor areas, the temporal, supramarginal and frontal gyrus, precuneus, anterior cingulate cortex, insula, and thalamus, among others, was associated with better episodic memory.^18^ In this study, higher variability in the cerebellum was associated with lower episodic memory performance.^18^ The authors suggested that increased variability was particularly beneficial for cognitive performance if it occurred in brain regions involved in integrating segregated functional domains.^18^ A two-time point rs-fMRI study of 74 healthy adults aged 19 to 82 years found that higher BOLD signal variability was associated with better working memory at both time points but was less robustly associated with episodic memory (only one time point showed this trend).^19^ Another study examined the relationship between variability and working memory performance in 20 healthy controls and 20 individuals at risk for Alzheimer’s disease, defined by a Montreal cognitive assessment (MoCA) score below 26 and reduced medial temporal lobe volume.^17^ In this study, greater voxel-wise variability, particularly in the medial temporal lobe, hippocampus, visual cortex and striatum, was associated with worse working memory performance in healthy controls but showed no relationship with working memory in the risk group.^17^ In contrast, previous findings appear to be more consistent for vocabulary performance, as three recent studies in adults found that brain signal variability during rest was not associated cross-sectionally with vocabulary.^18–20^ However, one study found that greater decline in brain signal variability over 2.5 years was associated with a more pronounced decline in vocabulary performance.^19^

Given that brain signal variability seems to play a role in both aging and cognition, it may also be a neural mechanism underlying resilient cognitive aging, i.e., cognitive reserve (CR). CR is a concept that aims to explain the disjuncture between the level of brain pathology and structural brain aging and cognitive performance. Individuals who have cognitive performance that is better than expected given their brain status are said to have high CR. Several factors have been identified that underlie CR, but the neural mechanisms require further research. Intra-individual variability in brain activity may be a rs-fMRI measure worth exploring as a neural mechanism underlying CR. To test for the presence of a true mechanism of CR, the CR-test proposed by the “Reserve and Resilience” Collaboratory (https://reserveandresilience.com/) should be used. This test states that neural mechanism of CR explains variance in cognitive performance beyond the variance explained by brain status and/or moderates the brain status-cognition relationship.^22,23^

The first aim of this project was to characterize associations between age and variability in brain activation during rest, as measured by rs-fMRI. Our second aim was to investigate whether intra-individual variability in brain activation during rest relates to cognitive performance in the domains of perceptual speed, episodic memory, fluid reasoning, and vocabulary. Lastly, we wanted to clarify whether brain signal variability during rest represents a neural mechanism underlying CR, by employing the formal test discussed above. We hypothesize that the brain signal variability changes with age and expect to find regional differences. In addition, based on the findings from previous studies,^15,17–21^ we hypothesize that greater brain signal variability is beneficial for perceptual speed, memory performance, and fluid reasoning, but does not play a role for vocabulary performance. Lastly, we hypothesize that brain signal variability represents a neural mechanism underlying CR.

## 2. Material and Methods

### 2.1. Participants

This project was conducted using data from the the Reference Ability Neural Network (RANN) and the Cognitive Reserve (CR) studies.^24–26^ These two studies, which are ongoing at Columbia University, include only participants who are free of psychiatric or medical conditions that could affect cognition. Resting-state fMRI data were available for 489 participants, with information on educational level available for all but three participants. At least one cognitive domain score could be calculated for 471 of these 486 participants with resting-state and complete covariate data. Of these 471 participants, we were able to calculate cognitive domain-specific brain status variables for all but one participant, leading to a final sample size of 470 participants.

### 2.2. Variability in brain activation during rest

Since we were interested in the variability in brain activation during rest, we first pre-processed the rs-fMRI data using a native spaced method developed in-house.^27^ In summary, we used FSL’s slice-timer tool for slice-timing correction and FSL’s mcflirt tool to register all volumes to a reference image, which was created by registering all volumes to the middle volume and averaging them using 6 df, 256 bins mutual information, and Sinc interpolation. Frame-wise displacement (FWD) was calculated according to Power et al.^28^ using the six motion parameters and the root mean square difference (RMSD) of the BOLD percentage signal in the consecutive volumes for each participant. RMSD was computed on the motion-corrected volumes before temporal filtering and we set the threshold of our RMSD to 0.3%, so that contaminated volumes could be detected and then replaced with new volumes generated by linear interpolation of adjacent volumes. The motion-corrected data were then filtered with a band-pass filter (cutoff frequencies of 0.01 and 0.09 Hz) using FSL’s fslmaths–bptf tool.^29^ In a last step, the motion-corrected, scrubbed, and temporally filtered volumes were residualized by regressing out the FWD, RMSD, lateral ventricular, and the left and right hemisphere white matter signals.^30^

To then quantify the degree of variability in brain activation over time, we calculated the standard deviation (SD) of the BOLD signal during the rs-fMRI scanning time for each brain region of interest (ROI) separately. Since the distributions of the temporal variability values were skewed, we log-transformed the SD values, also in line with the original version of the Scaled Subprofile Modelling (SSM) technique which assumed multiplicative variability and positive definite signal values. To then extract brain variability patterns, we applied the SSM principal component analysis (SSM-PCA) technique to the ROI-specific SD data. SSM is a multivariate technique with a long history of successful application and development in the Columbia group^31–33^ and can be used to derive BOLD variability-based brain patterns whose pattern scores maximally correlate with the outcome of interest. The SSM-PCA approach is based on the idea that functionally connected regions are activated together and that there are sources of variation that are shared by multiple brain regions. Thus, it performs PCA as a dimensionality reduction method and identifies several components that best capture the variability of regional brain activation. However, this method requires that there are no missing values for any ROI, so we first excluded all ROIs with missing values, resulting in a reduction from 264 to 223 ROIs.

To choose the optimal number of principal components (PCs) with respect to the cognitive outcome of interest, we selected all PCs with an eigenvalue greater than one and then used the Akaike Information Criterion (AIC) to select the number of PCs that gave the best model fit. We then used the optimal number of PCs to calculate the extent to which each person in the sample expresses the pattern of variability in each individual brain region as well as an overall pattern score. Afterwards, we created 500 random resamples (drawn with replacement) and determined the robustness of the pattern by calculating the inverse coefficient of variation for each ROI loading [z (loading) = point estimate / standard deviation (mean difference between point estimate and estimates from all resamples)]. The z-value roughly follows a standard normal distribution, so the z-value can be used to filter for those regions that are robustly associated with the cognitive outcomes (filtering for a z-value > 1.96 equals selecting those ROIs that are significantly associated at a p-value level of 0.05). To obtain the neuroanatomical labels of the significant ROIs, we mapped the coordinates of the regions from the Power atlas to the regions in the AAL atlas. We then visualized the brain regions with robust contributions to the pattern scores using BrainNet Viewer with the surface template “BrainMesh_CH2withCerebellum.nv”.^34^

### 2.3. Cognition

Cognitive performance was evaluated across four domains: perceptual speed, episodic memory, fluid reasoning, and vocabulary. Each domain was assessed with three cognitive tasks per domain during the neuropsychological examination. In addition, participants from the RANN study completed three computerized tasks per cognitive domain within the MRI scanner. As part of the neuropsychological examination, perceptual speed was assessed using the WAIS-III Digit-symbol task, the Stroop Color and Word Test,^35^ and the trail-making test A. In the MRI scanner, RANN participants performed the three perceptual speed tasks Digit Symbol, Letter Comparison, and Pattern Comparison tasks.^36^ To assess episodic memory performance outside the scanner, three measures from the Selective Reminding Test (SRT) were taken: long-term storage, continuous long-term retrieval, and number of words remembered in the last retrieval.^37^ Additionally, memory performance in the three in-scanner tasks Logical Memory, Word Order Recognition, and Paired Associates tasks was measured. To assess fluid reasoning performance, the neuropsychological examination included the Matrix Reasoning and the Letter-number Sequencing task of the Wechsler Adult Intelligence Scale III (WAIS-III) as well as the Block Design test. Within the scanner, we measured fluid reasoning with the Paper Folding, Matrix Reasoning and Letter Sets^38^ tasks. Lastly, vocabulary performance was measured with the WAIS-III Vocabular test, the Wechsler Test of Adult Reading,^39^ and the American National Adult Reading Test^40^ during the neuropsychological examination, and with the Synonyms,^41^ Antonyms,^41^ and Picture Naming tasks^42^ inside the scanner.

The assignment of cognitive tasks to cognitive domains was previously done using factor analysis, and we kept the same factor structure to be consistent with previous work. We then extracted each domain score using confirmatory factor analysis for all participants who had performance data available on at least two tasks in that domain (lavaan package in R^43^). Missing values were handled using full information maximum likelihood.

### 2.4. Characterization of brain status

T1-weighted magnetization-prepared rapid gradient echo (MPRAGE) scans were acquired in Tesla Philips Achieva Magnet scanners with a standard quadrature head coil using the following parameters: TE/TR of 3/6.5 ms, flip angle of 8°, in-plane resolution of 256 x 256, field of view of 25.4 × 25.4 cm, and 165–180 slices in axial direction with slice-thickness/gap of 1/0 mm. The FreeSurfer 5.1 analysis package was then used to reconstruct the structural T1 scan of each subject and to parcel the cortex into 68 ROIs according to the Desikan-Killiany cortical atlas. We then visually inspected the cortical parcellation and repeated the reconstruction in case of visual discrepancies until the results were satisfactory.

To calculate the mean cortical thickness for each of the 68 ROIs, we determined the distance between the gray/white matter surface and the gray/CSF surface at each point across the cortical mesh. To obtain the volume of the subcortical gray matter structures brain stem, hippocampus, amygdala, nucleus accumbens, nucleus caudatus, thalamus, putamen, and pallidum, we used the automatically segmented brain volume atlas (ASEG).^44^

To derive cognitive domain-specific brain status variables, we applied an approach that we have previously developed and successfully implemented.^45^ Specifically, we first regressed the mean cortical thickness of 68 ROIs, the volumes of eight subcortical gray matter measures, and ventricle sizes together on each of the four cognitive domain scores separately using generalized linear models with elastic net regularization (glmnet package in R^46^). We chose elastic net regularization because it effectively handles highly correlated predictor variables by selecting correlated variables together, rather than arbitrarily selecting one, while also removing those with no predictive value for the outcome.^47^ We applied 20-fold cross-validation and selected the model with the lowest cross-validated error for each cognitive domain. The selected model was then used to forward predict cognitive performance, and the resulting predicted values were extracted as cognitive domain-specific measures of brain status.

### 2.5. Characterization of cardiovascular risk

To characterize cardiovascular risk for each person, we constructed a variable that counted how many of the following self-reported risk factors were present for each person: diabetes, hypertension, history of heart disease, and smoking. For diabetes and hypertension, we also checked whether it was treated or untreated and coded no diabetes/hypertension as 0, treated diabetes/hypertension as 0.5, and untreated diabetes/hypertension as 1. We coded history of heart disease as 1 if any of the following questions were answered yes: myocardial infarction, congestive heart failure, other heart disease, peripheral vascular disease; otherwise, we coded it as 0. For smoking, we coded it as 1 if any of the following questions were answered yes: cigarette ever, cigarette now, cigar ever, cigar now, pipe ever, pipe now; otherwise, we coded it as 0. Thus, the cardiovascular risk score for each person could range between 0 and 1. For some participants we had data from two time points available and then selected the time point with the most complete data. We calculated a cardiovascular risk score if at least one risk factor was available.

### 2.6. Statistical analyses

To assess the associations between age and brain signal variability, we performed a multiple linear regression model with an age-related brain signal variability pattern score as outcome variable and age and sex as predictor variables.

Next, we assessed the association between brain signal variability and cognitive performance, by calculating one linear regression model for each cognitive domain score as outcome variable. We included a cognitive domain-specific brain signal variability pattern score as a predictor and additionally adjusted the model for age, sex and education.

To subject the brain signal variability pattern scores to the rigorous CR-test established by the “Reserve and Resilience” collaboratory,^22^ we calculated one regression model separately for each cognitive domain score as outcome variable that included the cognitive domain-specific brain status and brain signal pattern score as predictor variables. For brevity we just referred to “pattern scores” in the following remarks. The regression model reads as: *cognitive performance ∼ brain status + pattern scores + brain status*pattern scores + age + sex + education*. Age, sex, and education were included as covariates to account for performances differences that are not explained by differences in brain status or variability. A unique contribution by the variability value and/or a significant interaction term of brain status*pattern scores would fulfil the CR tenets.

Further, we conducted age- and sex-stratified regression analyses. For the age-stratified regression analysis, we divided the sample into two groups: participants under 60 years of age (N=237) and those aged 60 years or older (N=232). Within these age groups, as well as among women and men, we tested the associations between pattern scores and cognitive performance using the regression models described above. To assess whether these associations differed significantly between men and women and between younger and older participants, we drew 100 bootstrap resamples within each group. For each resample, we extracted the standardized regression coefficient for the pattern score within each stratum and computed all possible difference values between the two strata being compared, resulting in 10,000 beta difference values. We then checked the portion of beta difference values that deviated from most beta difference values by falling on the other side of zero. This inferential test allowed us to assess whether the association between the pattern score and cognitive performance differed significantly between men and women or between younger and older participants.

To account for confounding influences by differences in cardiovascular factors, we included a sum score for cardiovascular events in each regression model to see if the results change.

We performed all statistical analyses in R Studio (version 2022.12.0, R-base version 4.2.2).^48^ We corrected all our analyses for multiple testing applying the false discovery rate (FDR) correction (N=4 because of the four cognitive domains).

## 3. Results

### 3.1 Study sample

In the sample, 56.6% were women, the median age was 59 years (age range: 20 to 80 years) and the average educational level was high (mean=15.8 years, SD=2.3). Descriptive characteristics of the sample and the performance in the 24 cognitive tasks can be found in **Table 1**.

**Table 1.** Sample Characteristics.

| Characteristic | Participants | Missing values [%] |
| --- | --- | --- |
| **Sex**, N<sub>women</sub> (%) | 266 (56.6) | 0.0 |
| **Age**, median [IQR] in years | 59.0 [39.0, 67.0] | 0.0 |
| **Race**, N (%) |  | 0.0 |
| White | 290 (61.7) |  |
| Asian | 19 (4.0) |  |
| African American | 120 (25.5) |  |
| Native Hawaiian or other Pacific Islander | 7 (1.5) |  |
| Other | 34 (7.2) |  |
| **Education**, M (SD) in years | 15.8 (2.3) | 0.0 |
| **Cardiovascular risk score**, median [IQR] | 0.0 [0.0, 0.4] | 4.0 |
| **Perceptual speed** |  |  |
| WAIS-III Digit-symbol task [n correct in 90 s, max=93], M (SD) | 53.6 (14.0) | 1.7 |
| Stroop Color and Word Test [number of colors correctly said in 45 s], M (SD) | 71.6 (13.6) | 2.8 |
| Trail-making test A [completion time in s], median [IQR] | 25.0 [21.0, 33.0] | 2.1 |
| Digit Symbol [median RT in s across correct trials], M (SD) | 1.5 (0.2) | 27.9 |
| Letter Comparison [median RT in s across correct trials], M (SD) | 1.6 (0.3) | 28.3 |
| Pattern Comparison [median RT in s across correct trials], M (SD) | 1.5 (0.2) | 27.4 |
| **Episodic memory** |  |  |
| Selective Reminding Test (SRT): long-term storage [n words, max= 72], M (SD) | 44.5 (15.4) | 2.3 |
| SRT: continuous long-term retrieval [n words, max= 72], M (SD) | 34.5 (17.7) | 2.3 |
| SRT: last retrieval [n words recalled, max=12], median [IQR] | 10.0 [9.0, 12.0] | 2.3 |
| Logical Memory [% on time correct], median [IQR] | 75.0 [66.7, 85.0] | 26.0 |
| Word Order Recognition [% on time correct], M (SD) | 48.8 (22.6) | 26.0 |
| Paired Associates tasks [% on time correct], median [IQR] | 70.0 [50.0, 90.9] | 26.0 |
| **Fluid reasoning** |  |  |
| WAIS-III Matrix Reasoning [% on time correct], M (SD) | 64.7 (20.0) | 2.3 |
| WAIS-III Letter-number Sequencing task [% on time correct], M (SD) | 54.7 (15.2) | 2.3 |
| Block Design test [% on time correct], M (SD) | 58.8 (19.4) | 3.4 |
| Paper Folding [% on time correct], median [IQR] | 50.0 [30.8, 73.3] | 26.6 |
| Matrix Reasoning [% on time correct], median [IQR] | 45.5 [22.2, 66.7] | 25.3 |
| Letter Sets [% on time correct], median [IQR] | 75.0 [53.3, 86.7] | 26.4 |
| **Vocabulary** |  |  |
| WAIS-III Vocabular test [% correct], median [IQR] | 58.0 [49.0, 64.0] | 5.1 |
| Wechsler Test of Adult Reading [% correct], median [IQR] | 42.0 [33.0, 47.0] | 2.8 |
| American National Adult Reading Test [% errors], median [IQR] | 10.0 [5.0, 18.0] | 2.1 |
| Synonyms [% correct], M (SD) | 62.8 (22.6) | 28.7 |
| Antonyms [% correct], M (SD) | 56.5 (21.3) | 28.1 |
| Picture Naming task [% correct], M (SD) | 53.6 (18.4) | 31.5 |

*Note.* We indicated the mean (M) and interquartile range (IQR) for skewed variables and the mean and standard deviation (SD) for almost normally distributed variables. n = number, RT = reaction time, WAIS-III = Wechsler Adult Intelligence Scale III. The cardiovascular risk score is an average (range: 0 to 1) of four self-reported cardiovascular risk factors: diabetes, hypertension, history of heart disease, and smoking.

### 3.2 Age and brain signal variability

In the regression model including the age-related pattern score for brain signal variability as outcome variable, age as predictor, and sex as covariate, we found that older age was overall associated with a higher brain variability pattern score (standardized regression coefficient (ß)=0.372, 95% confidence interval (CI): 0.336 to 0.407, *p*<0.001). The brain region-specific analysis then showed that older age was robustly associated with higher brain signal variability in mainly subcortical and medial brain regions, but lower brain signal variability in the cortical brain regions (**Figure 1A**). The optimal number of PCs included in the age-related pattern score was 54.

**Figure 1.**
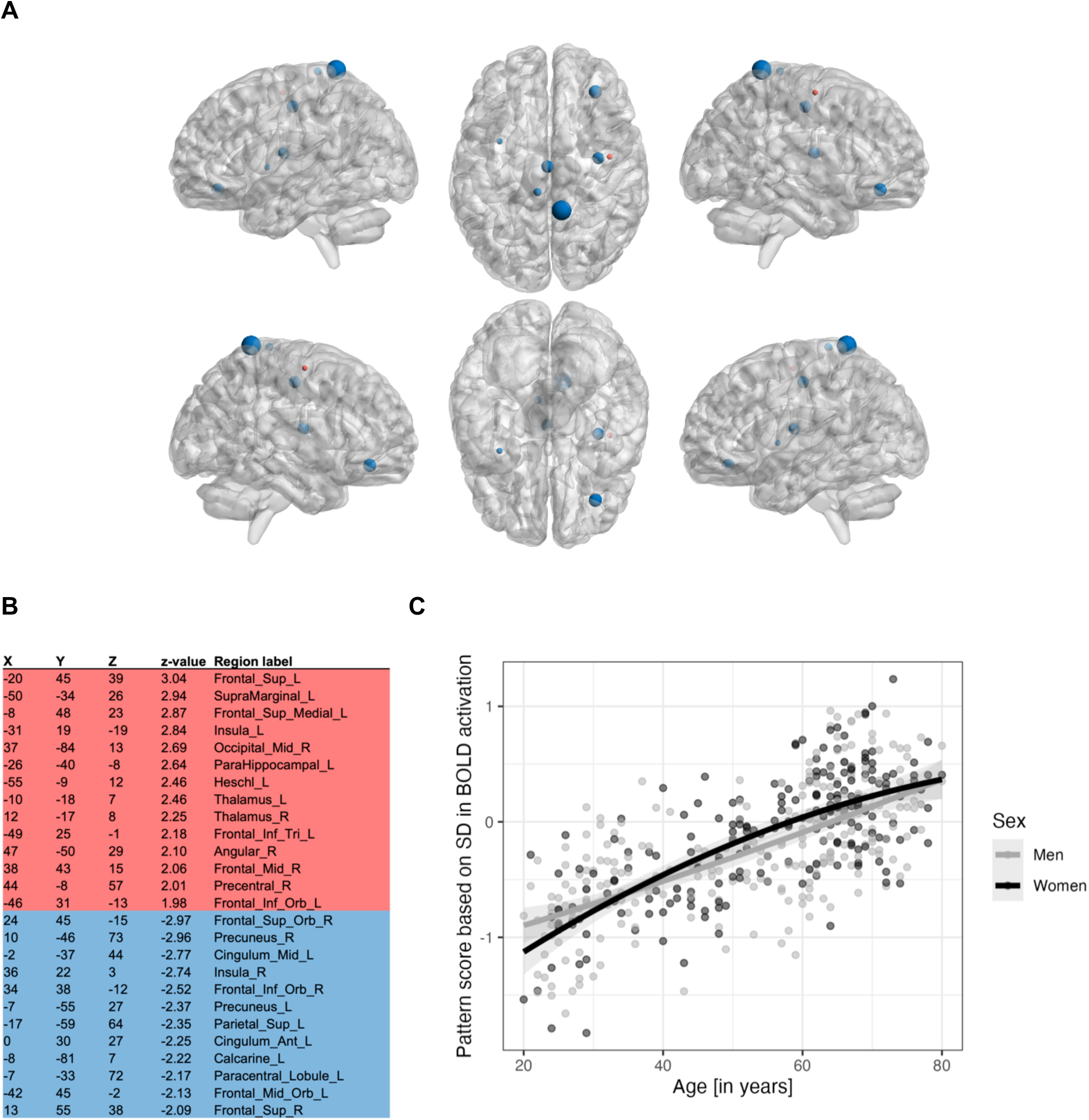
Age-related pattern scores for brain signal variability. **Panel A**: Visualization of cortical brain regions that had robust loadings in the age-related variability pattern score, as determined by a bootstrapping procedure. Red dots indicate cortical regions with positive loadings (higher expression of the age-related variability pattern score is associated with greater activation of these regions), and blue dots indicate cortical regions with negative loadings (higher expression of the age-related variability pattern score is associated with less activation of these regions). **Panel B:** List of brain regions that had robust loadings in the age-related variability pattern score, as determined by a bootstrapping procedure. If multiple coordinates of an AAL region robustly loaded on the pattern score, we selected the one with the highest z-value for the table and excluded those regions for which no AAL label could be assigned. The suffix “_L” denotes the left side, whereas “_R” denotes the right side. **Panel C**: Association between age [x-axis] and variability pattern score expression [y-axis], stratified by sex.

When we then stratified the analysis by sex, we found that women and men had similar expression of the pattern scores across the lifespan, as confidence intervals were largely overlapping (**Figure 1C**).

### 3.3 Brain signal variability and cognition

When we extracted cognitive domain-specific variability pattern scores, we found that the optimal number of PCs included in our pattern construction was 31 for perceptual speed, 22 for memory and reasoning, and 9 for vocabulary. Brain regions that significantly contributed to the brain signal variability pattern scores are shown in **Figure 2**. In the regression models that were adjusted for age, sex, and education, higher expression of the variability score was associated with faster perceptual speed (ß=0.276, 95% CI: 0.143 to 0.409, *p*<0.001, *p_FDR_*<0.001), better memory (ß=0.352, 95% CI: 0.207 to 0.498, *p*<0.001, *p_FDR_*<0.001), better reasoning (ß=0.400, 95% CI: 0.277 to 0.523, *p*<0.001, *p_FDR_*<0.001), and better vocabulary (ß=0.329, 95% CI: 0.224 to 0.435, *p*<0.001, *p_FDR_*<0.001) (**Figure 3A**). In the age-stratified regression analysis, we found similar results as in the main analysis for both age strata (**Figure 3B**). Using bootstrapping (N=100), we extracted a distribution of standardized regression coefficients for the association between pattern score and cognitive domains within each age stratum separately and then calculated all possible beta difference values between the two age groups (N=10,000 difference values). The mean beta difference was -0.147 for perceptual speed, -0.224 for episodic memory, -0.322 for fluid reasoning, and -0.133 for vocabulary, indicating that associations tended to be stronger in younger individuals. The bootstrap test indicated that the strength of the association between pattern score and reasoning differed significantly between young and old participants, as beta values were stronger in the younger group in 99.5% of bootstrap comparisons. For speed this was the case in 87.6% of comparisons, for memory in 93.6% of comparisons, and for vocabulary in 88.9% of comparisons.

**Figure 2.**
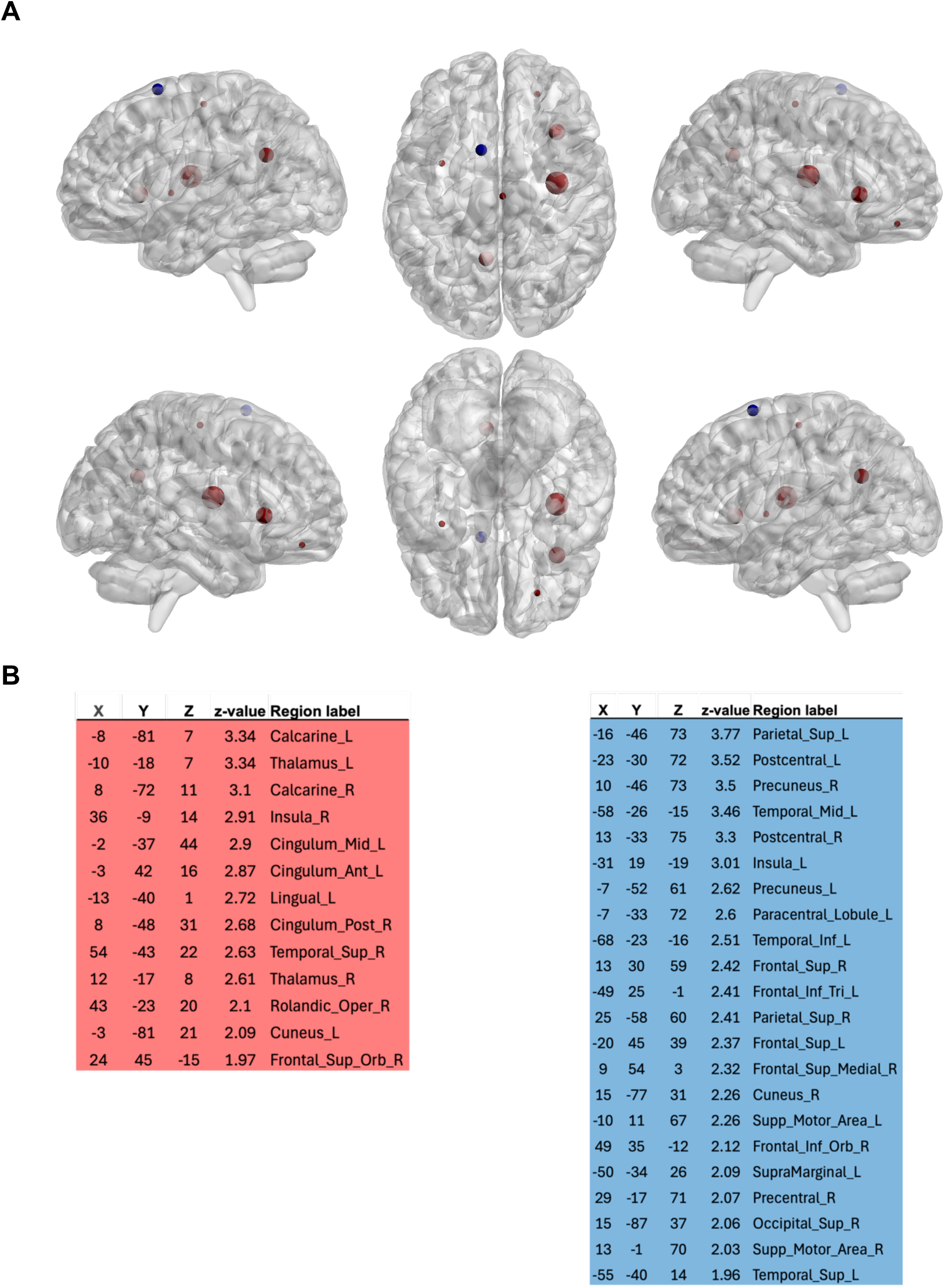
Brain regions with robust loadings in the cognition-related pattern scores for brain signal variability. **Panel A.** This graphic combines the results for the four cognitive domain-specific variability patterns in one graphic. If a node showed up more than once, we selected the node with the highest z-value. Red dots indicate cortical regions with positive loadings (higher expression of the cognition-related variability pattern score is associated with greater activation of these regions), and blue dots indicate cortical regions with negative loadings (higher expression of the cognition-related variability pattern score is associated with less activation of these regions). **Panel B.** The table shows all brain regions that had robust loadings in the cognition-related variability pattern scores, as determined by a bootstrapping procedure. If multiple coordinates of an AAL region robustly loaded on the pattern score, we selected the one with the highest z-value for the table and excluded those regions for which no AAL label could be assigned. In the table, the suffix “_L” denotes the left side, whereas “_R” denotes the right side.

**Figure 3.**
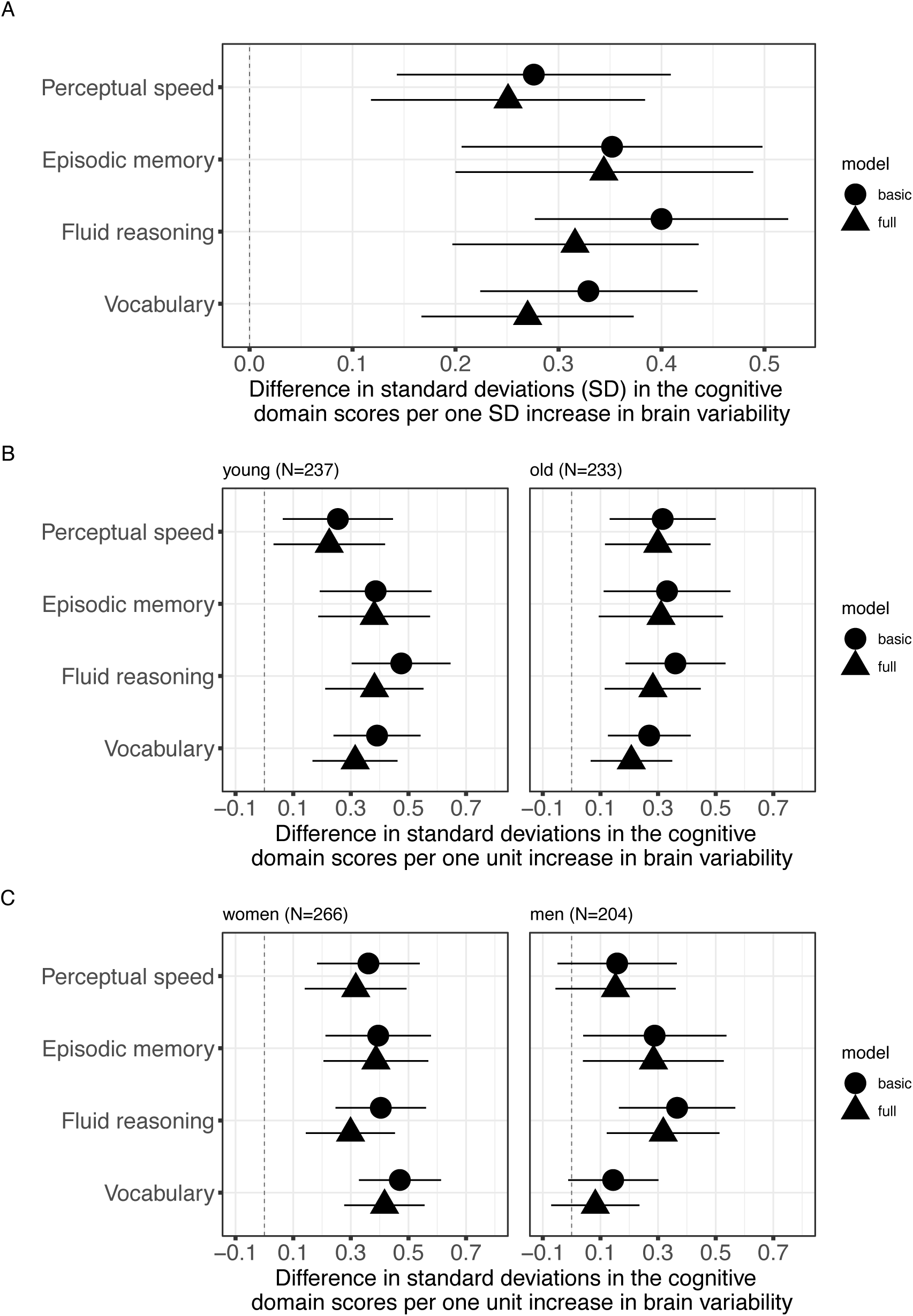
Associations between brain variability pattern scores and cognitive performance. The basic model includes the covariates age, sex, and education. The full model includes the covariates age, sex, education, and a cognitive domain-specific brain status variable. **Panel A**: Main results from the full sample. **Panel B**: Age-stratified analysis. **Panel C**: Sex-stratified analysis.

We then stratified all regression models by sex, using the cognitive domain-specific brain pattern scores that were derived in the full sample, and found that for women higher expression of the variability pattern was associated with faster perceptual speed (ß=0.361, 95% CI: 0.183 to 0.538, *p*<0.001, *p*_FDR_<0.001), with better episodic memory (ß=0.395, 95% CI: 0.212 to 0.578, *p*<0.001, *p*_FDR_<0.001), with better fluid reasoning (ß=0.403, 95% CI: 0.246 to 0.560, *p*<0.001, *p*_FDR_<0.001), and with better vocabulary (ß=0.470, 95% CI: 0.328 to 0.613, *p*<0.001, *p*_FDR_<0.001) (**Figure 3C**). In men, higher activation of the brain variability pattern scores was significantly associated with better fluid reasoning (ß=0.366, 95% CI: 0.164 to 0.567, *p*<0.001, *p*_FDR_<0.001) and better memory (ß=0.289, 95% CI: 0.040 to 0.537, *p*=0.023, *p*_FDR_=0.046), but not with performance in perceptual speed (ß=0.158, 95% CI: -0.049 to 0.365, *p*=0.135) or vocabulary (ß=0.144, 95% CI: -0.012 to 0.301, *p*=0.070) (**Figure 3C**). When we then compared the standardized regression coefficients for the associations between pattern scores and cognitive domain scores between men and women using bootstrapping, we found that the mean difference between men and women in standardized regression coefficients was -0.190 for perceptual speed, -0.092 for episodic memory, -0.036 for fluid reasoning, and -0.320 for vocabulary, indicating that associations between pattern scores and cognitive domains tended to be stronger in women. The results of the bootstrap test showed that women had stronger associations than men in 91.6% of the comparisons for perceptual speed, in 72.3% of the comparisons for episodic memory, in 60.1% of the comparisons for fluid reasoning, and in 99.9% of the comparisons for vocabulary. These results indicate that women showed significantly stronger associations between pattern scores and vocabulary than men.

### 3.4 Cognitive reserve tests

Adding a cognitive domain-specific brain status variable to the regression models that included a cognitive domain score as the outcome variable and age, sex, and education as covariates increased the variance explained by 1.0% in perceptual speed, 1.7% in episodic memory, 5.6% in fluid reasoning, and 5.1% in vocabulary. The regression results in the full sample suggest that higher variability pattern score is associated with faster perceptual speed beyond what would be expected given someone’s brain status, age, sex, and educational level (ß=0.250, 95% CI: 0.118 to 0.383, *p*<0.001, *p*_FDR_<0.001; **Figure 3A**). Similarly, the variability pattern score was associated with better memory performance (ß=0.345, 95% CI: 0.200 to 0.489, *p*<0.001, *p*_FDR_<0.001), better reasoning (ß=0.316, 95% CI: 0.196 to 0.436, *p*<0.001, *p*_FDR_<0.001), and better vocabulary (ß=0.270, 95% CI: 0.167 to 0.373, *p*<0.001, *p*_FDR_<0.001) even after adjusting for age, sex, education, and cognitive domain-specific brain status. We also found an interaction effect between brain status and brain variability pattern score expression on vocabulary (ß interaction term=-0.399, 95% CI: -0.723 to -0.075; *p*=0.016, *p*_FDR_=0.026) (**Figure 4**). All other interactions were not significant (*p*-values>0.677).

**Figure 4.**
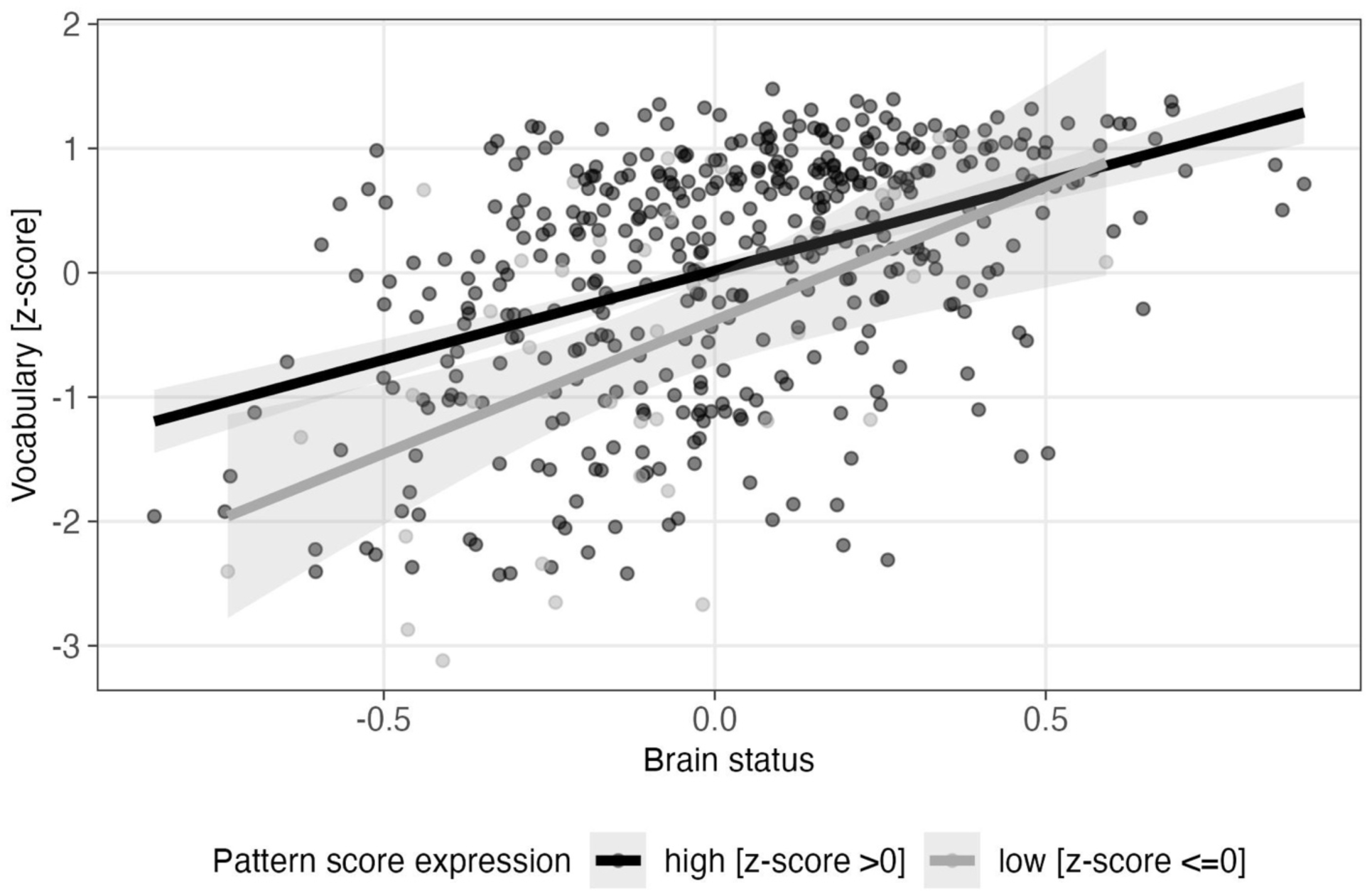
Interaction between brain variability pattern score for vocabulary and brain status on vocabulary performance. The scatterplot depicts on the x-axis the brain status variable that has been derived specifically for vocabulary performance. The y-axis depicts vocabulary performance. Each data point represents one participant. The black dots represent participants with a high expression of the brain variability pattern score (z-standardized score above 0) and the light-gray dots represent participants with low expression of the brain variability pattern score (z-standardized score below or equal to 0). For both groups of participants (high versus low expression of brain variability pattern score), there exists one superimposed function for the relationship between the brain status and cognitive change. The gray area around the group-specific regression lines signifies the 95% confidence interval for each group individually.

When we performed an age-stratified regression analysis with a split around 60 years, we found similar results to the main analysis (**Figure 3B**). The brain variability pattern scores did not modify the associations between brain status and the cognitive domain scores in the young and old age stratum (all *p*-values>0.107). The direction of the mean difference values of the standardized regression coefficients between the young and old age stratum that we obtained from bootstrapping indicated slightly stronger associations in the young compared to the older age stratum (-0.017 for perceptual speed, -0.171 for episodic memory, -0.195 for fluid reasoning, and -0.100 for vocabulary). However, the bootstrapping test was not significant for any of the comparisons, as the young group had stronger associations than the old group only in 56% of comparisons for speed, in 88% of comparisons for memory, in 94.2% of comparisons for reasoning, and in 82.8% of comparisons for vocabulary.

When we stratified the analysis for sex, we found that in women higher expression of the brain signal variability pattern scores was associated with faster perceptual speed (ß=0.316, 95% CI: 0.140 to 0.493; *p*<0.001, *p*_FDR_<0.001), with better episodic memory (ß=0.387, 95% CI: 0.205 to 0.569, *p*<0.001, *p*_FDR_<0.001), with better fluid reasoning (ß=0.298, 95% CI: 0.144 to 0.452, *p*<0.001, *p*_FDR_<0.001), and with better vocabulary performance (ß=0.416, 95% CI: 0.277 to 0.556, *p*<0.001, *p*_FDR_<0.001) (**Figure 3C**). The variability pattern scores did however not significantly modify the relationship between brain status and cognitive performance in women (all *p*-values > 0.054). In men, higher variability pattern scores were significantly associated with better fluid reasoning (ß=0.318, 95% CI: 0.122 to 0.513, *p*=0.002, *p*_FDR_=0.008) and better memory performance (ß=0.284, 95% CI: 0.039 to 0.529, *p*=0.023, *p*_FDR_=0.046), but not with perceptual speed (ß=0.152, 95% CI: -0.056 to 0.360, *p*=0.151) or vocabulary (ß=0.082, 95% CI: -0.071 to 0.235, *p*=0.291) after controlling for age, education, and cognitive domain-specific brain status (**Figure 3C**). There were also no significant moderation effects (all *p*-values > 0.587). The mean difference in standardized regression coefficients for the association between pattern score and cognitive domains between men and women was -0.151 for perceptual speed, -0.091 for episodic memory, 0.025 for fluid reasoning, and -0.325 for vocabulary, indicating that the association tended to stronger in women in all domains except reasoning. However, the bootstrapping test was only significant for vocabulary, where women had a higher regression coefficient for the association between pattern score and vocabulary than men in 99.9% of the comparisons.

### 3.5 Sensitivity analysis – cardiovascular risk

The results for the association between age and the variability pattern scores did not change when we additionally included the cardiovascular risk score in the regression model. Similarly, the results for the association between variability pattern scores and cognition did not change for the full sample and the age strata after including this additional covariate. The results of the sensitivity analysis for the CR test were also comparable for the full sample and for old age strata. However, the interaction effect between brain status and variability pattern score expression on vocabulary, that we found in the main analysis of the full sample, was no longer significant when the cardiovascular risk score was included in the model (ß interaction term = - 0.304, 95% CI: -0.638 to -0.030; *p*=0.074). In the young age group, the CR test for perceptual speed was no longer met when the cardiovascular risk score was included in the model (ß=0.196, 95% CI: -0.009 to 0.400, *p*=0.060). In the sex-stratified sensitivity analysis, the variability pattern scores were no longer significantly associated with memory performance in men (ß = 0.227, 95% CI: -0.030 to 0.485; *p*=0.083). Similarly, the CR test for memory did not remain significant after correcting for multiple testing when the additional covariate was included (ß = 0.282, 95% CI: 0.032 to 0.532; *p*=0.027, *p*_FDR_=0.054).

## 4. Discussion

In this project we first analysed whether the variability in BOLD signal during rest differs between adults aged 20 to 80 years, and whether it is associated with cognition. We then assessed whether it represents a neural mechanism underlying CR. We found region-specific differences in how age relates to brain signal variability, in line with our hypothesis. For the associations with cognition and CR in different domains, we hypothesized that brain signal variability would be relevant for perceptual speed, memory performance, and fluid reasoning, but not for vocabulary. However, the results from the main analysis suggest that brain signal variability plays a role for all four cognitive domains and fulfills the CR test for all four domains. Our stratified analyses showed that when assessing the association between brain signal variability and cognition, the association was significantly stronger in younger individuals for reasoning. In addition, the association between brain signal variability and vocabulary performance was significantly stronger in women than in men for both the cognition and the CR test.

We found that there are strong regional differences in how age relates to brain signal variability, with BOLD signal variability showing an age-related increase in regions including the left superior frontal gyrus, the left supramarginal gyrus, the insula, the left parahippocampal gyrus, and the thalamus. An age-related increase in BOLD signal variability in the superior frontal gyrus has also been reported in recent rs-fMRI studies.^4,14,15,18^ There is also some overlap between our study and previous fMRI studies regarding brain regions where older age was associated with lower variability. Older age was associated with lower activation of structures that belong to the default mode network (e.g., precuneus and cingulum) and the calcarine gyrus, in line with findings from previous studies.^4,15^ Despite the complexity of the results, the visualization in **Figure 1A** indicates a general tendency for variability to increase with age in subcortical and medial brain regions (only one red dot is visualized using the surface-based template), and to decrease with age in cortical regions, such as the default mode network or the primary visual cortex. Previous studies have also reported a general pattern of higher age-related variability in subcortical structures.^4,15^

Since greater brain signal variability goes along with more exploration of available functional network configurations, higher brain signal variability may imply greater flexibility of the brain.^2,5–7^ This could lead to more optimal responses to environmental demands.^2,5^ However, it has also been suggested that lower BOLD signal variability allows for better signal transmission.^8,9^ Thus, there may be an optimal level of brain signal variability, and once variability is higher than this optimum, variability may be detrimental, leading to reduced signal transmission and lower performance.^49^ The optimum level may vary depending on the function of a brain region (i.e., sensory signal transmission versus association areas). To better understand the observed age-related increases in brain signal variability, its associations with behavioural outcomes are important.

Regarding behavioural relevance of brain signal variability, we found that the results were very similar between the model adjusted for age, sex and education, and the fully adjusted model that we used to test whether brain signal variability underlies CR (**Figure 3**). Therefore, we will focus our discussion on the full model in the following. In the full sample, higher variability scores were associated with faster perceptual speed, and better episodic memory, fluid reasoning and vocabulary performance than would be expected given someone’s brain status, age, sex, and educational level. This means that the brain signal variability met the rigorous CR test proposed by the “Reserve and Resilience” Collaboratory, as BOLD signal variability explains variance in cognitive performance over and above the variance explained by brain status.^22,23^ We also observed that brain signal variability moderated the association between brain status and vocabulary performance and the visualization of the interaction effects indicates that in those individuals with higher expression of the vocabulary pattern score, vocabulary performance was less strongly dependent of brain status compared to those individuals with lower pattern score expression (Figure 4). When we subsequently age-stratified our analysis, we found the same associations in both age strata and a trend towards stronger associations in the young group (<60 years). However, the two age groups differed significantly only for fluid reasoning and only in the cognition model, not in the fully adjusted CR model. This suggests that brain signal variability represents an important neural mechanism of CR across the adult life span but has stronger behavioural consequences for fluid reasoning in younger age. We also showed that many brain areas are involved (**Figure 2)**.

One possible reason why we found a stronger association between brain signal variability and fluid reasoning in the young compared to the older subsample could be that age leads to suboptimal levels of brain signal variability. We found that those regions in which higher variability was beneficial for cognitive performance party overlap with those regions in which variability was lower in older age and the other way around. For example, lower variability (i.e., negative loadings in the variability pattern) in the superior frontal gyrus turned out to be included in the variability patterns that relates to CR, but this region shows higher variability in older age. And for the cingulum, where higher variability seems to benefit CR, we observed an age-related decrease in variability. Greater brain signal variability in frontal and subcortical brain regions may serve a compensatory mechanism in older adults to counteract age-related brain changes, though it does not necessarily translate into improved cognitive performance. There is already some evidence that the level of brain signal variability is indeed influenced by the status of the brain, as a study of 28 healthy adults (mean age at baseline: 72 years) showed that changes in cortical thickness partly accounted for age-related changes in brain signal variability.^50^ Further, greater white matter integrity, as measured by global fractional anisotropy, has been associated with greater resting-state brain signal variability after accounting for age in a study including 66 healthy adults aged 60 to 80 years.^18^

When we stratified our analyses by sex, we observed that higher brain signal variability was associated with better performance in all cognitive domains in women, but only with better episodic memory and fluid reasoning in men. The bootstrapping test then showed that the strength of the association between brain signal variability pattern score and cognition differed significantly between men and women only for vocabulary, with women showing a significantly stronger association. One possible reason may be that women, with a mean age of 52.5 years, were about two years younger than men (mean age: 54.9 years). As the age-stratified analysis showed that associations tended to be stronger in younger individuals (although not statistically significant for vocabulary), unmeasured factors associated with differences in the age distribution between women and men may partly explain the observed sex differences. However, further research in other large resting-state fMRI samples is needed to investigate possible sex differences in more detail.

We also conducted a sensitivity analysis in which we additionally controlled for cardiovascular risk to ensure that our results are not confounded by cardiovascular factors. Our results showed that the associations were largely unaffected by cardiovascular factors. However, we created a cardiovascular risk factor based on externally measured vascular risk factors but ideally, we would have controlled for cardiovascular factors by checking the BOLD response to hypercapnia.^51^ Previous research indicates that brain signal variability is a relatively robust marker,^51^ but we cannot completely exclude the possibility that our results are affected by differences in cardiovascular factors.

We chose rs-fMRI as data from rs-fMRI may be more generalizable than data from task-related fMRI studies, which are cognitive task-specific. In addition, rs-fMRI and task-related signal amplitude fluctuations have been shown to be linearly related across subjects, which suggests common underlying neuronal and physiological mechanisms.^52^ Still, it would be interesting to check if similar results can be achieved in samples having both rs-fMRI and task-related fMRI data available. One factor that limits the generalisability of the results is that our sample was on average highly educated, which likely limits the range of CR levels.

## 5. Conclusions

The level of BOLD signal variability showed differences as a function of age, with higher values with age for subcortical and medial regions, but lower values for cortical regions. Brain signal variability is associated with cognitive performance and represents a neural mechanism underlying CR across the adult lifespan. Sex differences require further investigation in other large samples, as we found that brain signal variability showed much stronger associations with vocabulary performance in women compared to women.

## 6. Acknowledgements

We would like to thank all participants from the Reference Ability Neural Network Study and the Cognitive Reserve Study. Many thanks also to the staff members of these studies for their contributions to data collection and data management.

## 8. Data availability

All data examined in the manuscript are available in pseudonymized format upon request.

